# EVIDENCE FOR ANGIOTENSIN II AS A NATURALLY EXISTING SUPPRESSOR FOR THE NATRIURETIC PEPTIDE SYSTEM

**DOI:** 10.1101/2023.01.26.525806

**Authors:** Xiao Ma, Seethalakshmi R. Iyer, Xiaoyu Ma, Shawn H. Reginauld, Yang Chen, Shuchong Pan, Ye Zheng, Dante Moroni, Yue Yu, Lianwen Zhang, Valentina Cannone, Horng H. Chen, Carlos M. Ferrario, S. Jeson Sangaralingham, John C. Burnett

## Abstract

**Background:** Natriuretic peptide system (NPS) and renin angiotensin aldosterone system (RAAS) function oppositely at multiple levels. While it has long been suspected that angiotensin II (ANGII) may directly suppress NPS activity, no clear evidence to date support this notion.

**Objectives:** This study was designed to systematically investigate ANGII-NPS interaction in humans, in vivo, and in vitro for translational insights.

**Methods:** Circulating atrial, b-type, and c-type natriuretic peptides (ANP, BNP, CNP), cyclic guanosine monophosphate (cGMP), and ANGII were simultaneously investigated in 128 human subjects. Prompted hypothesis was validated in rat model to determine influence of ANGII on ANP actions. Multiple engineered HEK293 cells and surface plasmon resonance (SPR) technology were leveraged for mechanistic exploration.

**Results:** In humans, ANGII showed inverse relationship with ANP, BNP, and cGMP. In regression models predicting cGMP, adding ANGII levels and interaction term between ANGII and natriuretic peptide increased predicting accuracy of base models constructed with either ANP or BNP, but not CNP. Importantly, stratified correlation analysis further revealed positive association between cGMP with ANP or BNP only in subjects with low, but not high, ANGII levels. In rats, co-infusion of ANGII even at physiological dose attenuated blood pressure reduction and cGMP generation triggered by ANP infusion. In vitro, we showed that the suppression effect of ANGII on ANP-stimulated cGMP requires the presence of ANGII type-1 (AT_1_) receptor and mechanistically involves protein kinase C (PKC), which can be substantially rescued by either valsartan (AT_1_ blocker) or Go6983 (PKC inhibitor). Using SPR, we showed ANGII has low affinity for particulate guanylyl cyclase A (GC-A) receptor binding compared to ANP or BNP.

**Conclusions:** Our study reveals ANGII as a natural suppressor for cGMP-generating action of GC-A via AT_1_/PKC dependent manner and highlights importance of dual-targeting RAAS and NPS in maximizing beneficial properties of natriuretic peptides in cardiovascular disease.

**STRUCTURED GRAPHICAL ABSTRACT:** 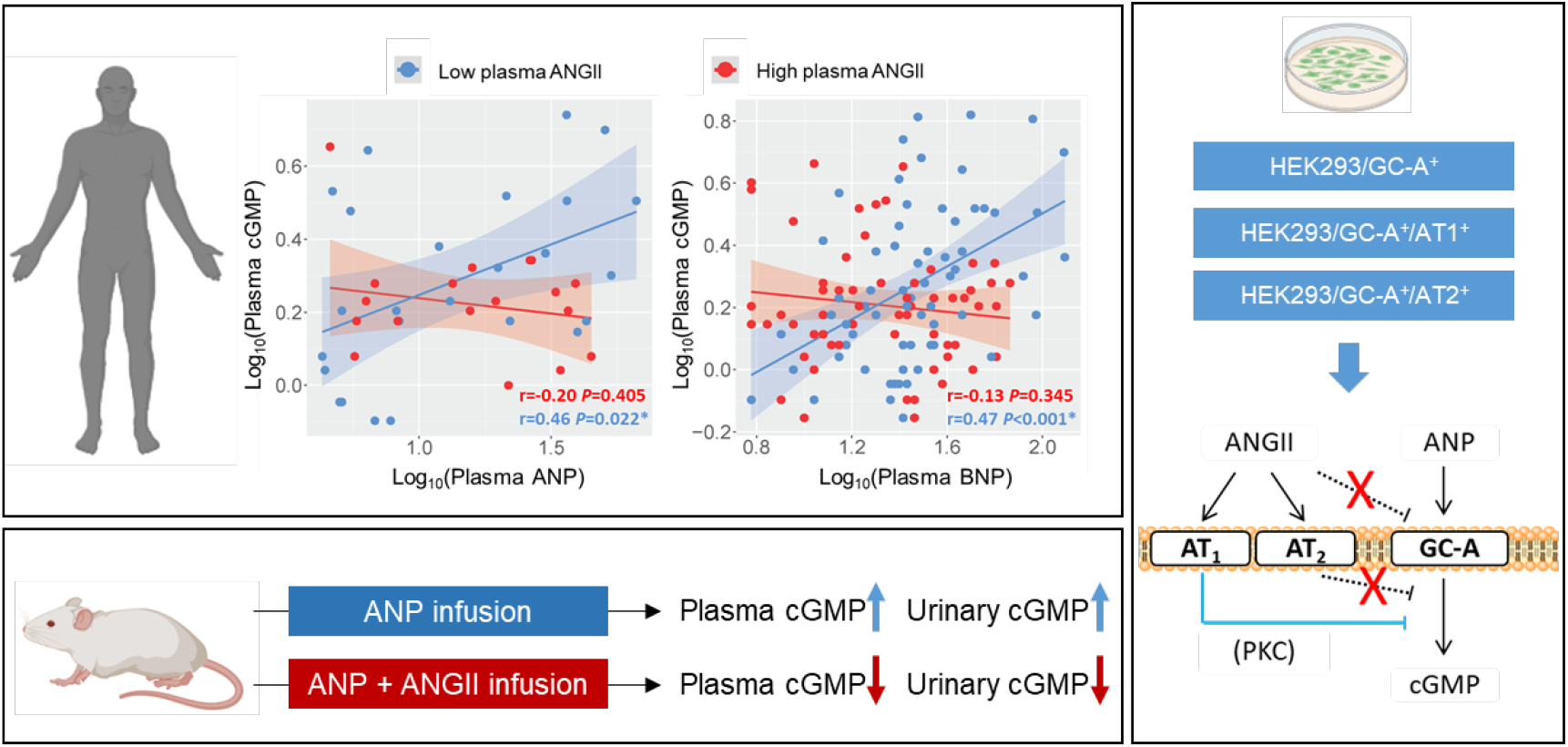

**CLINICAL PERSPECTIVES:** Accumulating evidence continues to support the NPS as a promising therapeutic target via the function of the GC-A receptor and production of the second messenger cGMP for heart failure, hypertension, and other cardiovascular diseases. Improving our mechanistic understanding on GC-A/cGMP pathway regulation may further advance the development of novel NPS enhancing therapies. Here we report evidence from multiple avenues supporting a fundamental, yet previously underappreciated mechanism involving a negative action of ANGII in suppressing GC-A receptor-mediated cGMP production via an AT_1_ receptor-dependent manner. This study also provides a solid rationale for the superiority of combinatory neurohormonal therapies such as sacubitril/valsartan in treating cardiovascular disease, and further highlights a promising therapeutic avenue of dual targeting both the NPS and RAAS to maximize protection.

## INTRODUCTION

The natriuretic peptide system (NPS) serves a fundamental role in controlling blood pressure (BP), volume homeostasis, and the functional and structural adaptation of the heart and kidneys to physiologic or pathophysiologic stresses (1-6). To date, the family of human natriuretic peptides (NPs) includes the cardiac-derived hormones atrial natriuretic peptide (ANP) and b-type natriuretic peptide (BNP), and the renovascular-derived hormone c-type natriuretic peptide (CNP) (7-11). In response to hypertension-induced cardiovascular remodeling or diseases such as heart failure (HF), diabetes, and chronic kidney disease (CKD), production of all three peptides can up-regulate to exert pluripotent beneficial actions including vasodilatation, diuresis, natriuresis, prevention of myocardial hypertrophy, and suppression of cardiac and renal fibrosis. As such, the augmentation of endogenous NPs under pathophysiologic conditions is considered as compensatory, and strategies to further enhance their local and circulating levels, by either providing exogenous analogs (12) or inhibiting peptide degradation (13), are attractive for therapeutic development.

At the molecular level, the functional receptor for ANP and BNP is particulate guanylyl cyclase A (GC-A, also known as NPR1 or NPRA), while GC-B (also known as NPR2 or NPRB) is the molecular target for CNP (3,7,14-16). Upon NP binding, both GC-A and GC-B undergo conformational changes to generate its second messenger 3’, 5’ cyclic guanosine monophosphate (cGMP) for cardiorenal protection. However, there has been limited mechanistic understanding on how the activity of these receptors are regulated, which may hinder the development of optimal NP-based therapeutics.

A conventional therapeutic strategy for treating cardiovascular (CV) disease is the targeting of angiotensin II (ANGII), the central hormone of the renin angiotensin aldosterone system (RAAS) (17-19). It is well established that the deleterious action of RAAS under pathophysiologic conditions is mediated via the ANGII type 1 (AT_1_) receptor, and drugs blocking ANGII generation or activity, such as angiotensin converting enzyme (ACE) inhibitors and AT_1_ receptor blockers (ARBs), remain first-line therapies for CV disease (20-23). Further, a well-recognized interaction exists between RAAS and the NPS (24,25). Along the line, a series of recent clinical trials have established the effectiveness of sacubitril/valsartan (S/V) in the clinical management of patients with HF (26-28) and hypertension (29), further underscoring the therapecutic value of this imporant crosstalk between RAAS and NPS as sacubitril prevents ANP and BNP degradation via inhibiting neprilysin (NEP) while valsartan blocks ANGII from binding to AT_1_ receptor. However, studies in the past have disproportionally focused on how activation of the NPS counteracts RAAS activity. For instance, a mouse model with global knock-out of GC-A gene (*Npr1*) exhibits enhanced cardiac expression of RAAS components including ACE and AT_1_ receptor (30,31). Comparatively, far fewer investigations have been performed to understand if any of the endogenous RAAS components influence the beneficial actions of the NPS from a therapeutic and pharmacological perspective. In fact, ANGII has been reported to antagonize the activity of NPS via inhibiting ANP-induced cGMP under multiple in vitro conditions (32-34), but whether and how ANGII may influence the activity of NPS in vivo and in humans remains uncertain.

Here our goal was to systematically investigate the influence of ANGII on the NPS with a focus on GC-A. We simultaneously assessed circulating ANGII, ANP, BNP, CNP, and cGMP as well as defined their correlation and interaction in 128 healthy subjects to understand this crosstalk under physiologic conditions. The findings from the human study were expanded using a rodent model of ANGII and ANP infusion in vivo. Additionally, we interrogated the molecular mechanisms of the interaction between ANGII and the NPS with multiple engineered cells in vitro. Together, we provide strong evidence in support of the concept that ANG II serves as a naturally existing suppressor of the NPS by influencing the activity of GC-A receptor and cGMP generation.

## METHODS

### Study Population

128 healthy subjects (71% are female) were recruited from the Mayo Clinic Biobank (MCB) as we have previously described (11). The MCB is a comprehensive and ongoing registry for collecting biological samples including blood from volunteers to aid ongoing population studies which focus on precision medicine, medical diagnostics, and therapeutics. All subjects enrolled in current study have been confirmed to be non-smokers, with no history of CV or systemic diseases and were not taking any CV medications at the time of sample collection. This study was approved by the Institutional Review Board (IRB) at Mayo Clinic and all participants provided written informed consent for participation. All plasma samples were collected on enrollment and stored at -80ºC until further assayed and analysis.

### Biomarker Measurements in Human Plasma

Plasma ANP, BNP, CNP, ANGII, and cGMP were measured in the Mayo Clinic Cardiorenal Research Laboratory. Plasma ANP was measured with a radioimmunoassay (RIA) developed at the Mayo Clinic (Phoenix Pharmaceuticals, Burlingame, CA) (35). The standard curve range of the ANP assay was from 4.8 to 1,250 pg/mL with a minimum detectable value at 4.0 pg/mL. The cross-reactivity of the ANP assay is at < 1% using BNP, CNP, ANGII. Plasma BNP was measured with a 2-site immunoenzymatic sandwich assay (Biosite Inc, Alere, France). The BNP concentrations were determined from a stored multipoint calibration curve with a range between 5 and 4,000 pg/mL. There was no cross-reactivity with ANP or CNP in this assay. Plasma CNP was measured with a non-equilibrium radioimmunoassay kit (Phoenix Pharmaceutical, Burlingame, CA) using an antibody against human CNP as previously reported (11). The range of the standard curve used in the assay was 0.5 to 128 pg, with the lowest detection of 0.5 pg. There was no detectable cross-reactivity with ANP, urodilatin and BNP below 600 pg/mL, and the cross-reactivity with BNP at 600 pg/mL was 1.3%. Plasma ANGII was measured with a polyclonal rabbit antibody (Phoenix Pharmaceuticals, Burlingame, CA) with a minimum level of detection of 0.5 pg/tube. The inter- and intra-assay variations were 13% and 9%. The cross-reactivity of ANGII assay is 100% with ANGIII and [Val^5^]-ANGII, 0.9% with angiotensinogen (AGT), 0.5% with ANGI, 0.0% with ANP or BNP. Plasma cGMP was measured by ELISA (Enzo Life Sciences, Farmingdale, NY) following the manufacturer’s instructions.

Concentrations of plasma creatinine were measured by the Clinical Chemistry Core Laboratory at the Mayo Clinic, using a Cobas creatinine reagent (Roche Diagnostics, Indianapolis, Indiana). The plasma creatinine values were used to calculate estimated glomerular filtration rate (eGFR) using the chronic kidney disease epidemiology collaboration (CKD-EPI) equation.

### Synthetic Peptides

Throughout the in vivo and invitro study, human ANP 1-28 (catalog#005-06, Phoenix Pharmaceuticals, Burlingame, CA), human BNP 1-32 (catalog#011-03, Phoenix Pharmaceuticals, Burlingame, CA), and human ANGII (catalog#A9525, Sigma-Aldrich, St. Louis, MO) were used.

### In Vivo Rat Experiments

Protocols in this study have been approved by the Mayo Clinic Institutional Animal Care and Use Committee (IACUC). Rats (Sprague-Dawley, male, 10-11 weeks old) were anesthetized and maintained with 2% isoflurane. Rats were kept on heating pad at 37 ºC for the entire experiment to maintain their body temperature. Polyethylene tubes (PE-50) were placed into a jugular vein for saline and peptides infusion and into a carotid artery for BP monitoring and blood sampling. The bladder was cannulated with a PE-90 tube for passive urine collection. After the surgical setup, an initial infusion of 0.9% saline started at a fixed rate based on rat weight and allow to equilibrate for 45 mins. After the 45-min equilibrium, BP was recorded, and blood was sampled as baseline reference values. After another 5-min of recovery, infusion of saline control or peptides (ANP, ANGII, or ANP+ANGII) was initiated, and urine collection was also initiated. Based on our preliminary testing, ANP at 300 pmol/kg/min and ANGII at 50 pmol/kg/min were chosen for the current protocol. A blood sample was collected from the carotid artery at 15, 30, 60, and 90 mins post initiation of peptides (or saline control) infusion and placed in EDTA tubes on ice. Urine samples were collected every 30 mins period and BP was measured every 15 mins using CardioSOFT Pro software (Sonometrics Corporation). Blood was centrifuged at 2500 rpm at 4 ºC for 10 mins, and the plasma was aliquoted. The volume of urine was measured, and the sodium concentrations were measured with an electrocyte analyzer (Diamond Diagnostics Inc, Holliston, MA). Both plasma and urine samples were then stored at -80 ºC until assayed. cGMP was measured in rat plasma and urine samples by a cGMP ELISA (Enzo Life Sciences, Farmingdale, NY) as previously reported (36).

### Generation and Maintenance of HEK293 Transfected Cell Lines

HEK293 cells were maintained in Dulbecco’s modified Eagle’s medium (DMEM) supplemented with 10% fetal bovine serum, 100 U/mL penicillin, 100 U/mL streptomycin, and designated anti-biotics. HEK293/GC-A^+^ and HEK293/GC-B^+^ cell lines were generated from HEK293 parental cell transfected with plasmids (OriGene, Rockville, MD) containing either human GC-A or GC-B cDNA sequences. Both GC-A and GC-B plasmid carried sequences for a green florescence protein (GFP) tag and resistance to Neomycin (G418). HEK293/GC-A^+^/AT_1_^+^ and HEK293/GC-A^+^/AT_2_^+^ cell lines were generated from HEK293/GC-A^+^ transfected with lentiviral particle with clones of either human AT_1_ or AT_2_ receptor (OriGene, Rockville, MD) using polybrene transfection agent. Initial transfection was done with multiplicity of infection (MOI) at 20, 10, and 1. The lentiviral vector carried sequence for resistance to puromycin. Positive HEK293/GC-A^+^ and HEK293/GC-B^+^ cells were cultured in regular medium with 250 μg/mL G418; Likewise, positive HEK293/GC-A^+^/AT_1_^+^ or HEK293/GC-A^+^/AT_2_^+^ cells were cultured in regular medium with 250 μg/mL G418 (to select for GC-A^+^ cells) and 1 μg/mL puromycin (to select for AT_1_^+^ or AT_2_^+^ cells).

### Intracellular cGMP Generation in HEK293 Transfected Cell Lines and human primary renal cells

HEK293/GC-A^+^, HEK293/GC-A^+^/AT_1_^+^, HEK293/GC-A^+^/AT_2_^+^ cells were plated in a 24-well plate and cultured to reach 80-90% confluency before treatment. The treatment buffer included 0.5 mM 3-isobutyl-1-methylxanthine (IBMX, Sigma-Aldrich, St. Louis, MO) to prevent cGMP degradation. Cells were the treated with treatment buffer (vehicle), 10^−8^, 10^−7^, or 10^−6^ M ANP with and without 10^−8^ M ANGII for 10 mins before being harvested. Cells were then lysed, sonicated, centrifuged, and the supernatants were extracted and reconstituted in 300 μL 0.1M HCl for cGMP assay. The samples were then assayed using a cGMP ELISA (Enzo Life Sciences, Farmingdale, NY). To study the effect of valsartan or Go6983 (protein kinase C inhibitor) on cGMP, 10^−6^ M valsartan (Sigma-Aldrich, St.Louis, MO) or 5 μM Go6983 (Bio-Techne Corporation, Minneapolis, MN) was also added together with treatment buffer and peptides as described above.

Human renal proximal tubular epithelial cells (HRPTCs) (ScienCell Research Laboratories, Carlsbad, CA) were maintained and subcultured in according to the manufacturer’s protocols. For cGMP generation by ANP with or without ANGII, HRPTCs were cultured in 6-well plate and same procedure were followed as in HEK293 cells.

### Membrane and Cytosol Fractionation on HEK293 Transfected Cell Lines

HEK293/GC-A^+^ and HEK293/GC-A^+^/AT_1_^+^ cells were plated in a 100mm petri dishes and cultured to reach 80-90% confluency before treatment of 10^−8^M ANGII for 10 mins. Cells were then collected in cold PBS and sujected for membrane and cytosol separation using Mem-PER Plus Membrane Protein Extraction Kit (Thermo Fisher Scientific, Waltham, MA). Both GC-A and NaK-ATPase, two known cell membrane proteins, were used as the control for a successful separation of the membrane fraction from the cytostol fraction.

### Western Blotting

To collect whole cell lysis, HEK293 parental cells, HEK293/GC-A^+^, HEK293/GC-B^+^, HEK293/GC-A^+^/AT_1_^+^, HEK293/GC-A^+^/AT_2_^+^ cells were cultured in 60 mm dish to reach 90-100% confluency before being harvested in RIPA buffer (Thermo Fisher Scientific, Waltham, MA) with protease inhibitors. Alternatively, cells were cultured and prepared following procedure described above to separate the membrane and cytosol fractions. The samples were then incubated for 30 mins, centrifuged, and collected for the supernatant. Western blotting was performed with standard protocol and 40 μg of protein from each sample were used for blotting. The following primary antibodies were used: anti-GCA (catalog#MAB48601, R&D System, Minneapolis, MN) at 1:1000 dilution, anti-GCB (catalog#55113-1-AP, Proteintech, Rosemont, IL) at 1:200 dilution, anti-GFP (catalog#TA150041, OriGene, Rockville, MD) at 1:1000 dilution, anti-PKCα (catalog#2056, Cell Signaling Technology, Danvers, MA) at 1:500 dilution, anti-PKCε (catalog#2683, Cell Signaling Technology, Danvers, MA) at 1:1000 dilution, anti-Phspho-(Ser) PKC Substrate (catalog#2261, Cell Signaling Technology, Danvers, MA) at 1:1000 dilution, anti-NaK-ATPase (catalog#EP1845Y, abcam, Waltham, MA) at 1:2000 dilution, anti-p38 MAPK (catalog#8690, Cell Signaling Technology, Danvers, MA) at 1:1000 dilution, anti-Phospho-p38 MAPK (catalog#4511, Cell Signaling Technology, Danvers, MA) at 1:1000 dilution, and anti-GAPDH (catalog#2118, Cell Signaling Technology, Danvers, MA) at 1:5000 dilution.

### Real-time Quantitative PCR

HEK293/GC-A^+^, HEK293/GC-A^+^/AT_1_^+^, HEK293/GC-A^+^/AT_2_^+^ cells were cultured in 60 mm dish to reach 90-100% confluency before being harvested. Four biological replicates were conducted. mRNA was extracted using TRIzol (Thermo Fisher Scientific, Waltham, MA) following the manufacturer’s instruction. cDNA was synthesized by using Superscript III First-Strand Synthesis System (Thermo Fisher Scientific, Waltham, MA) using 1 μg mRNA. Real-time reverse transcription PCR assays were conducted in 96-well plates using the LightCycler 480 Instrument (Roche, Wilmington, MA). Levels of mRNA expression were normalized to glyceraldehyde 3-phosphate dehydrogenase (*GAPDH*). The primers sequences for angiotensin II receptor type 1 (*AGTR1*) gene are 5-’ATTTAGCACTGGCTGACTTATGC-3’ and 5’-CAGCGGTATTCCATAGCTGTG-3’. The primers sequences for angiotensin II receptor type 2 (*AGTR2*) gene are 5’-AAACCGGTTCCAACAGAAGC-3’ and 5’-GAGCCTCAAAGCAAGTAGCC-3’.

### GC-A Binding Studies

Surface plasmon resonance (SPR) measurements were performed on a BI-4500 SPR instrument (Biosensing Instrument Inc. Tempe AZ) at 25°C as previously described (37). Briefly, 40 ug/ml of extracellular domain human GC-A recombinant protein (MyBioSource, San Diego, CA) containing 12 histidine residues on the C-terminus was immobilized to the linked nickel sulfate on the Ni-NTA sensor chip (Biosensing Instrument, Tempe, AZ). The chip was then washed with buffer (150 mM NaCl, 50 μM EDTA, pH 7.4, 0.1% DMSO), and sequentially diluted ANGII, ANP or BNP were injected at the rate of 60 μL/min and allowed to dissociate for 200 seconds. Binding kinetics were derived from sensorgrams using BI-Data Analysis Program (Biosensing Instrument, Tempe, AZ). Affinity analysis of GC-A with ANGII, ANP, and BNP were performed using a 1:1 Langmuir binding model. Two series were performed for all studies.

### Statistical Analysis

For human data, categorical variables were presented as count and percentage. Continuous variables which followed a normal distribution were presented as mean ± SD. Continuous variables which did not follow an approximate normal distribution were presented as median (IQR), and values were log transformed to approximate normality for further regression analyses. Univariable regression models were constructed to determine independent characteristics associated with the log transformed ANGII and other variables, and the Pearson correlation coefficient (r) was presented together with the associated *P* value for each model to indicate the trend of either positive or negative association. In multiple regression models for cGMP prediction, the absolute value as well as change in coefficient of determination (R^2^) and the associated *P* values were used as the primary metric for model evaluation. Data analysis and visualization were conducted with R programing language (version 4.1.1).

All in vitro and in vivo data were presented as mean ± SEM or median (interquartile range [IQR]). Two-tailed unpaired t test or non-parametric Mann-Whitney test were used for two groups comparison. One-way ANOVA followed by Dunnett post hoc test was performed for multiple group comparison. Two-way ANOVA with repeated measures followed by Sidak post hoc test was performed in studies with multiple time points involved. Data analysis and visualization were conducted in GraphPad Prism 9 (GraphPad Software, La Jolla, CA), and statistical significance was accepted as *P* < 0.05.

## RESULTS

### Study Population

The baseline characteristics of the 128 healthy subjects are presented in **Table 1**. Under healthy conditions, the circulating levels of NPs, cGMP, and ANGII were low. There was no significant difference in baseline characteristics between male and female subjects (**Supplemental Table 1**). Approximately 65% of healthy subjects (N = 84) had plasma ANP levels at or below the minimal detectable value of 4 pg/mL, and these subjects were excluded for ANP-related regression analyses described below.

**Table 1.**
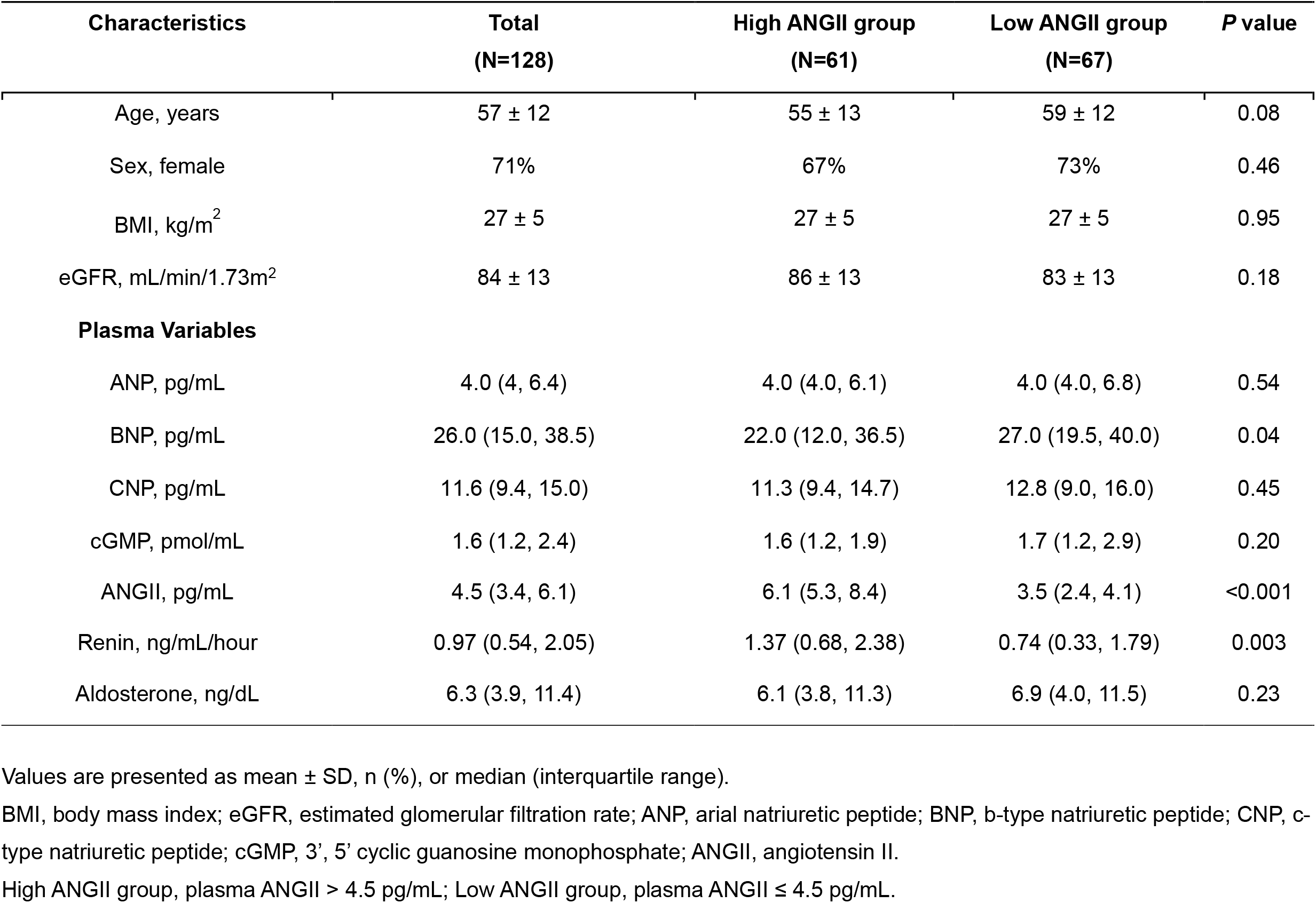
Baseline Characteristics of Healthy Subjects.

| Characteristics | Total<br>(N=128) | High ANGII group<br>(N=61) | Low ANGII group<br>(N=67) | P value |
| --- | --- | --- | --- | --- |
| Age, years | 57 ± 12 | 55 ± 13 | 59 ± 12 | 0.08 |
| Sex, female | 71% | 67% | 73% | 0.46 |
| BMI, kg/m<sup>2</sup> | 27 ± 5 | 27 ± 5 | 27 ± 5 | 0.95 |
| eGFR, mL/min/1.73m<sup>2</sup> | 84 ± 13 | 86 ± 13 | 83 ± 13 | 0.18 |
| **Plasma Variables** |  |  |  |  |
| ANP, pg/mL | 4.0 (4, 6.4) | 4.0 (4.0, 6.1) | 4.0 (4.0, 6.8) | 0.54 |
| BNP, pg/mL | 26.0 (15.0, 38.5) | 22.0 (12.0, 36.5) | 27.0 (19.5, 40.0) | 0.04 |
| CNP, pg/mL | 11.6 (9.4, 15.0) | 11.3 (9.4, 14.7) | 12.8 (9.0, 16.0) | 0.45 |
| cGMP, pmol/mL | 1.6 (1.2, 2.4) | 1.6 (1.2, 1.9) | 1.7 (1.2, 2.9) | 0.20 |
| ANGII, pg/mL | 4.5 (3.4, 6.1) | 6.1 (5.3, 8.4) | 3.5 (2.4, 4.1) | <0.001 |
| Renin, ng/mL/hour | 0.97 (0.54, 2.05) | 1.37 (0.68, 2.38) | 0.74 (0.33, 1.79) | 0.003 |
| Aldosterone, ng/dL | 6.3 (3.9, 11.4) | 6.1 (3.8, 11.3) | 6.9 (4.0, 11.5) | 0.23 |

Values are presented as mean ± SD, n (%), or median (interquartile range).
BMI, body mass index; eGFR, estimated glomerular filtration rate; ANP, arial natriuretic peptide; BNP, b-type natriuretic peptide; CNP, c-type natriuretic peptide; cGMP, 3', 5' cyclic guanosine monophosphate; ANGII, angiotensin II.
High ANGII group, plasma ANGII > 4.5 pg/mL; Low ANGII group, plasma ANGII ≤ 4.5 pg/mL.

### Correlations between ANGII and Natriuretic Peptides in Humans

Median plasma ANGII levels was 4.5 (IQR 3.4 - 6.1) pg/mL. Univariable analysis (**Supplemental Table 2**) revealed that levels of ANGII tend to decrease with age (r = - 0.13, *P* = 0.15), and there was an inverse relationship between ANGII with ANP (r = - 0.24, *P* = 0.12) or BNP (r = -0.16, *P* = 0.07), but not CNP (r = -0.02, *P* = 0.81). Importantly, plasma cGMP and ANGII were also weakly associated in a negative manner (r = -0.14, *P* = 0.11).

**Table 2.** Regression Models to Predict Plasma cGMP in Healthy Subjects.

| Model | R<sup>2</sup> | ΔR<sup>2</sup> | P<sub>model</sub> | P<sub>interaction</sub> |
| --- | --- | --- | --- | --- |
| ANP base model<sup>#</sup> | 0.275 | --- | 0.039 | --- |
| ANP base model<sup>#</sup> + ANGII + ANP*ANGII | 0.330 | 0.055 | 0.028 | 0.06 |
| BNP base model | 0.133 | --- | 0.006 | --- |
| BNP base model + ANGII + BNP*ANGII | 0.225 | 0.092 | <0.001 | <0.001 |
| CNP base model | 0.143 | --- | 0.009 | --- |
| CNP base model + ANGII + CNP*ANGII | 0.166 | 0.014 | 0.012 | 0.20 |

ANP base model includes age, sex, BMI, eGFR, and ANP; BNP base model includes age, sex, BMI, eGFR, and BNP; CNP base model includes age, sex, BMI, eGFR, and CNP.
ANP\*ANGII, BNP\*ANGII, and CNP\*ANGII refer to the interaction terms between ANGII with ANP, BNP or CNP.
<sup>#</sup>N = 44 as subjects with plasma ANP at or below the minimal detection level (4 pg/mL) were excluded for this analysis.

### ANGII Interacts with the Correlations between cGMP and ANP/BNP in Humans

At the molecular level, ANP, BNP, and CNP all stimulate cGMP production via GC receptors. To further elucidate the potential influence of ANGII on the actions of NPs, we constructed a series of regression models to predict plasma cGMP (**Table 2**). As expected, a base model (with age, sex, BMI, and eGFR) that contained ANP, BNP or CNP alone showed significant predictive value for cGMP (R^2^ > 0.13, *P* < 0.05 for all). Surprisingly, we found that further adding ANGII levels and the interaction term between ANGII and the corresponding NP into base model enhanced the accuracy in cGMP prediction dramatically with ANP (ΔR^2^ = 0.055, *P*_*interaction*_ = 0.06) or BNP (ΔR^2^ = 0.092, *P*_*interaction*_ < 0.001), but less with CNP (ΔR^2^ = 0.014, *P*_*interaction*_ = 0.20).

We then stratified these healthy subjects by the median level of ANGII (4.5 pg/mL) and investigated the correlations between cGMP and each NP separately. Comparison in baseline characteristics between high and low ANGII groups are shown in **Table 1**. Interestingly and as shown in **Figure 1**, plasma cGMP was found to be significantly and positively associated with ANP (r = 0.46, *P* = 0.022) or BNP (r = 0.47, *P* < 0.001) only in those with lower ANGII (≤ 4.5 pg/mL), but not in those with a higher ANGII (> 4.5 pg/mL). By contrast, no significant association was found between cGMP and CNP in either ANGII group **(Supplemental Figure 1)**. Thus, our data on healthy subjects raise a possibility that ANGII can disrupt the “connection” between the NPs and cGMP even under physiologic condition and this interaction appears to be more towards GC-A (the receptor for ANP and BNP) rather than GC-B (the receptor for CNP).

**Figure 1.**
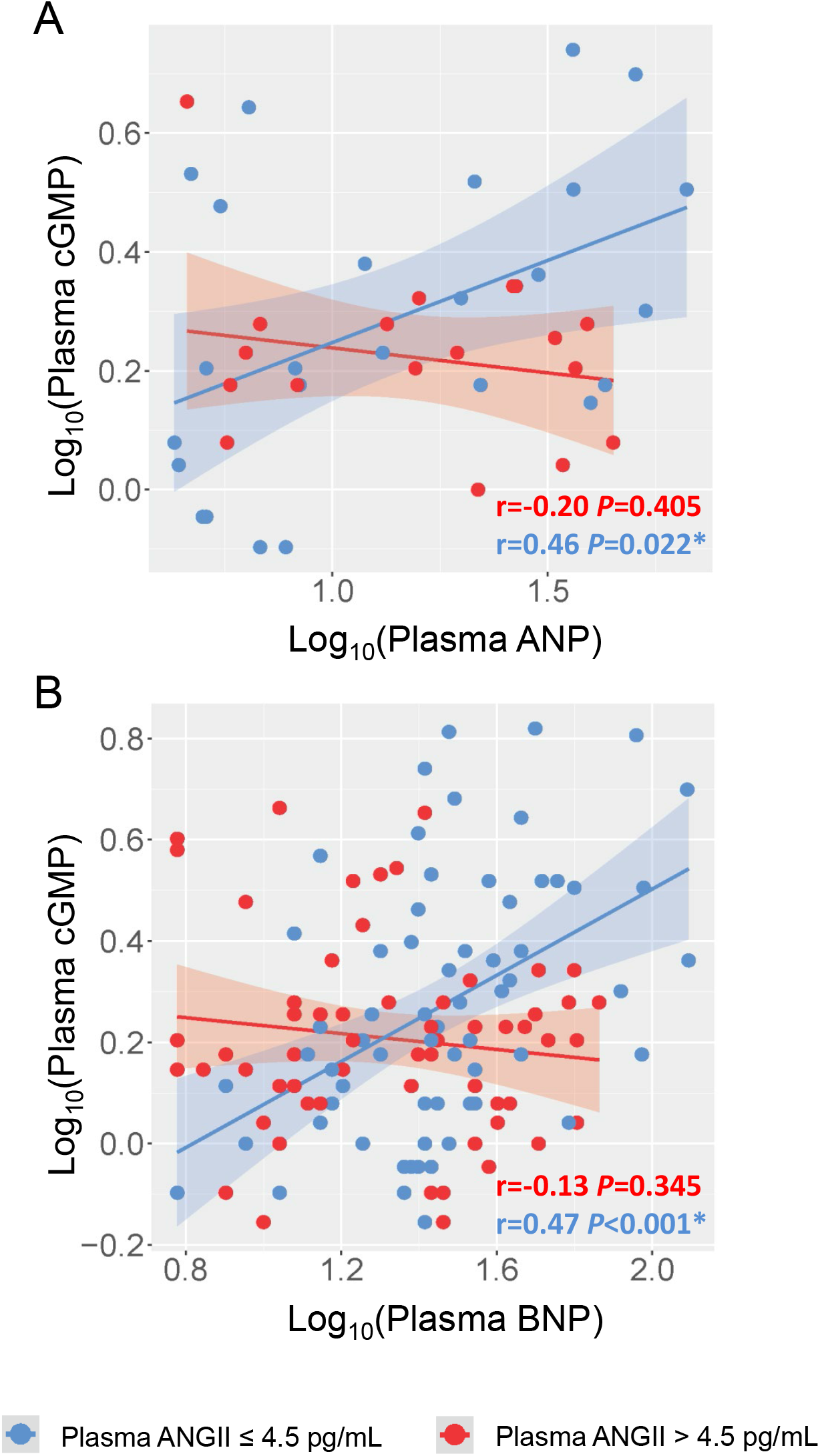
Correlations between cGMP and ANP/BNP in Healthy Subjects Grouped by ANGII Levels. (A) Linear correlation between plasma ANP and plasma cGMP. (B) Linear correlation between plasma BNP and plasma cGMP. Each dot represents one healthy subject. Blue indicates subjects with plasma ANGII ≤ 4.5 pg/mL; Red indicates subjects with plasma ANGII > 4.5 pg/mL. In (A), subjects with plasma ANP below the minimal detection by the assay (4 pg/mL) was not included. Pearson correlation coefficient r and associated *P* value were shown for each constructed linear regression model. * indicates *P* < 0.05.

### ANGII Attenuates ANP-induced cGMP Production In Vivo

To validate the insights gained from our human study, we investigated the influence of ANGII on GC-A activity in vivo by infusing ANP, which is a more potent generator of cGMP than BNP (15,38), in the presence or absence of ANGII in the normal rats. We first determined a physiological dose of ANGII via infusing different doses of ANGII alone into normal rats (**Supplemental Figure 2** and data not shown). Continuous infusion of ANGII at 50 pmol/kg/min resulted in a mild increase in BP, no increase in urine flow, and no change in circulating ANP levels during the 90-min infusion period (**Supplemental Figure 2A-2D**). More importantly, infusion of ANGII (50 pmol/kg/min) had no effect on plasma and urinary cGMP (**Supplemental Figure 2E-2F**) at any time points during the infusion. Therefore, we leveraged ANGII at this dose for the co-infusion experiments with ANP (**Figure 2A**).

**Figure 2.**
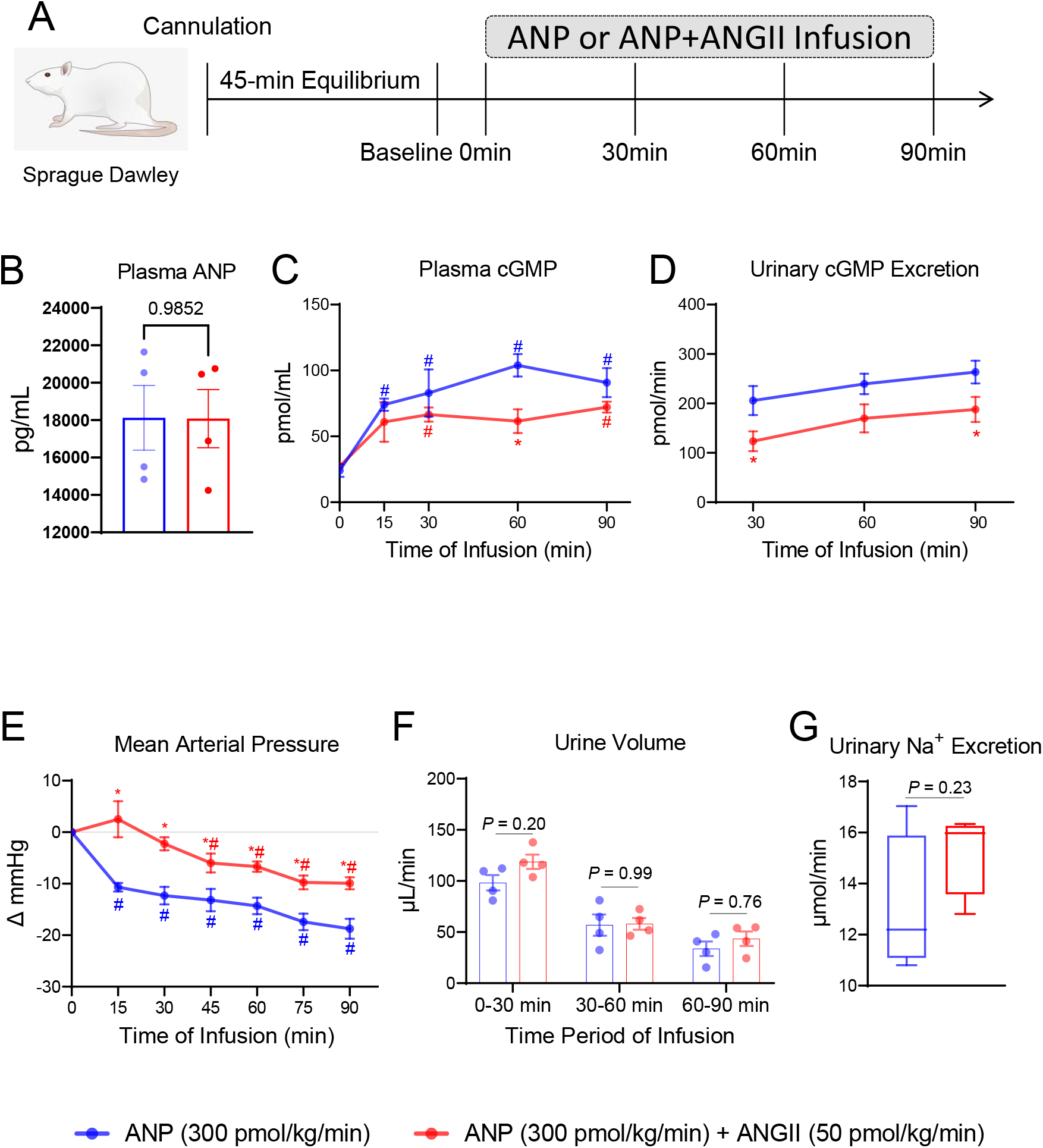
ANGII Attenuates Circulating and Urinary cGMP Generation Induced by ANP In Vivo. (A) The acute protocol study design to evaluate the effect of ANGII on ANP infusion in normal rats. (B) Plasma ANP levels at 90 mins after continuous ANP infusion. (C) Plasma cGMP, * indicates *P*<0.05 compared to ANP group; # indicates *P*<0.05 compared to corresponding baseline. (D) Urinary cGMP excretion. * indicates *P*<0.05 compared to ANP group. (E) Change in mean arterial pressure. * indicates *P*<0.05 compared to ANP group; # indicates *P*<0.05 compared to corresponding baseline. (F) Urine output represented by urine volume collected at every 30-min interval during continuous infusion. (G) Urinary sodium excretion during the entire 90-min continuous infusion. N=4 in ANP group, N=4 in ANP+ANGII group.

Continuous infusion of ANP at 300 pmol/kg/min alone increased plasma and urinary cGMP levels and reduced BP throughout the 90-min infusion period in normal rats (**Figure 2C-2E**). While the endogenous circulating ANP levels was determined to be 15.3±1.4 pg/mL, infusion of ANP at this dose increased circulating ANP levels to 18125.1±1729.3 pg/mL (**Figure 2B, Supplemental Figure 2D**). Compared to vehicle, ANP infusion (300 pmol/kg/min) also enhanced urine volume throughout (**Figure 2F, Supplemental Figure 2C**). By contrast, co-infusion of ANGII (50 pmol/kg/min) together with ANP (300 pmol/kg/min) decreased plasma cGMP consistently and reached statistical significance (*P* = 0.019, two-way ANOVA followed by multiple comparison) at 60 mins (**Figure 2C**). Meanwhile, urinary cGMP was consistently lower in rats infused with both ANP and ANGII compared to those infused with ANP alone, and there were significant differences at 30 mins (206 vs 123 pmol/min, *P* = 0.036) and 90 mins (263 vs 188 pmol/min, *P* = 0.042) (**Figure 2D**). There was also an attenuation on ANP-induced BP lowering effect by ANGII (**Figure 2E**). Meanwhile, we observed no influence of ANGII on the urine volume or sodium excretion that incurred with ANP infusion (**Figure 2F-2G**). Of note, we confirmed that the co-infusion of ANGII had no influence on circulating ANP levels (**Figure 2B**).

### ANGII Suppresses GC-A Activity via AT_1_ Receptor

There are two well-recognized functional receptors for ANGII in humans. While AT_1_ receptor is thought to mediate most of the deleterious actions by ANGII, the function of ANGII type 2 (AT_2_) receptor is generally thought to have salutary actions (39). To further gain mechanistic insights into how ANGII may suppress GC-A mediated cGMP production, we engineered 3 different HEK293 cell lines which included GC-A overexpression (HEK293/GC-A^+^), GC-A and AT_1_ co-overexpression (HEK293/GC-A^+^/AT_1_^+^), and GC-A and AT_2_ co-overexpression (HEK293/GC-A^+^/AT_2_^+^) **(Figure 3A**). Compared to parental HEK293 cells which has negligible endogenous GC-A expression, all three engineered cell lines had substantial GC-A overexpression as confirmed at the protein level (**Figure 3B**). Given the challenges with specific antibodies against ANGII receptors (40,41), we validated AT_1_ and AT_2_ overexpression with quantitative PCR (**Figure 3C**). Compared to HEK293/GC-A^+^, the mRNA expression of *AGTR1* (human AT_1_ coding gene) was up-regulated by ∼429 folds in HEK293/GC-A^+^/AT_1_^+^ cells and the mRNA expression of *AGTR2* (human AT_2_ coding gene) was up-regulated by ∼5 folds in HEK293/GC-A^+^/AT_2_^+^ cells, both indicating the successful overexpression of functional receptors of ANGII.

**Figure 3.**
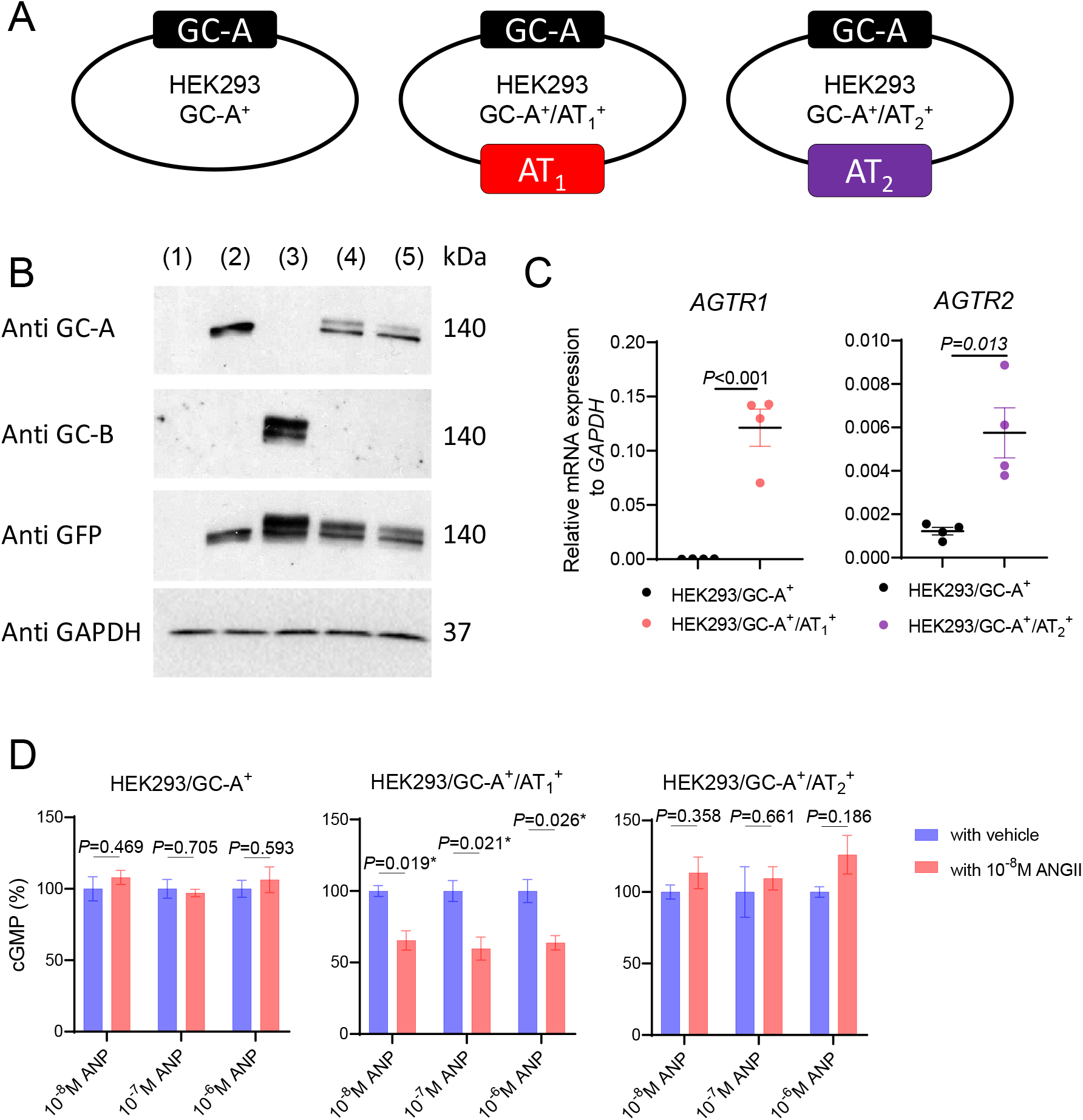
ANGII Attenuates GC-A Mediated cGMP Generation via AT_1_ Receptor. (A) Construction and nomenclature of three different HEK293 transfected cells. (B) Protein expression of human GC-A and GC-B in HEK293 parental and transfected cells by western blotting. 1, HEK293 parental cells; 2, HEK293/GC-A^+^; 3, HEK293/GC-B^+^; 4, HEK293/GC-A^+^/AT_1_^+^; 5, HEK293/GC-A^+^/AT_2_^+^. Each lane was loaded with 40 μg total protein. Antibody against GFP detected protein expression incurred by either GC-A or GC-B transfection. HEK293/GC-B^+^ cells served as a negative control for GC-A specific overexpression in other transfected cells. (C) mRNA expression of human *AGTR1* (AT_1_ coding gene) in HEK293/GC-A^+^/AT_1_^+^ cells and human *AGTR2* (AT_2_ coding gene) in HEK293/GC-A^+^/AT^2+^ cells, compared to HEK293/GC-A^+^ cells. Expression levels were normalized to human *GAPDH*. (D) In vitro cGMP generation in HEK293 transfected cells in response to different doses of ANP with or without ANGII (10^−8^ M). Values of cGMP in ANGII treated group (red) were normalized to corresponding vehicle group (blue) under each dose of ANP. Absolute values of cGMP are shown in **Supplemental Figure 3**. * indicates *P*<0.05, two-way ANOVA with Sidak multiple comparisons test. N=3 biological replicates (defined as cells grown in 3 independent plates) in each designed group.

To investigate if AT_1_ receptor or AT_2_ receptor may be involved in the crosstalk between ANGII and GC-A, we stimulated the 3 engineered cell lines with ANP alone or in the presence of ANGII (**Figure 3D**). Interestingly, while ANP alone dose-dependently increased cGMP production in all three transfected cell lines (**Supplemental Figure 3**), the presence of ANGII significantly reduced the ANP-induced cGMP production only in HEK293/GC-A^+^/AT_1_^+^ cell (by ∼40% reduction), but not in HEK293/GC-A^+^ or HEK293/GC-A^+^/AT_2_^+^ cells (**Figure 3D**). Therefore, we conclude that the presence of AT_1_ receptor, but not AT_2_ receptor, is required for the ANGII-associated suppression on GC-A receptor. Besides HEK293 cells, we also recapitulated a dose-depedent suppression effect of ANGII on ANP-induced cGMP in human renal proximal tubular epithelial cells (HRPTCs), a primary cells expressing most of the NPS and RAAS membrane receptors including GC-A, AT_1_ and AT_2_ (**Supplemental Figure 4**).

**Figure 4.**
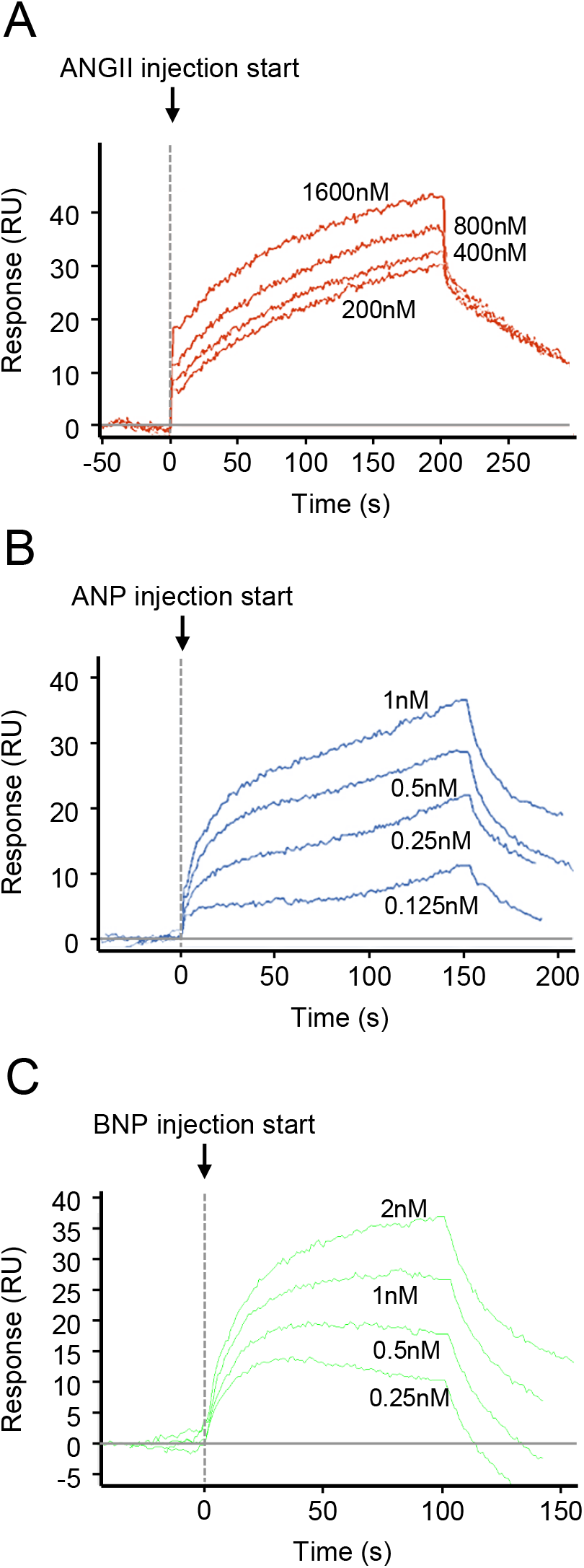
SPR Binding Curves for ANGII, ANP, and BNP to Human GC-A receptor. Representative of SPR sensogram for the binding of ANGII (A), ANP (B), and BNP (C) to the extracellular domain of human GC-A receptor.

### Binding Kinetics of ANGII to Extracellular Domain of Human GC-A receptor

We also determined the binding kinetics of ANGII to GC-A. Surface plasmon resonance (SPR) analysis was conducted for increasing concentration of ANGII to the extracellular domain of human GC-A receptor (**Figure 4A**). The affinity constant K_D_ of the interaction between ANGII and GC-A was found to be 258 nM (**Table 3**). As a reference, strong binding between either ANP or BNP to human GC-A was validated with K_D_ of 0.342 nM and 0.843 nM, respectively (**Figure 4B-4C, Table 3**). Compared to ANP and BNP, the binding between ANGII and GC-A was also associated with both a low association constant (K_a_) and a low dissociation constant (K_d_) (**Table 3**), which altogether suggests that ANGII may have low, if any, binding affinity to human GC-A receptor.

**Table 3.**
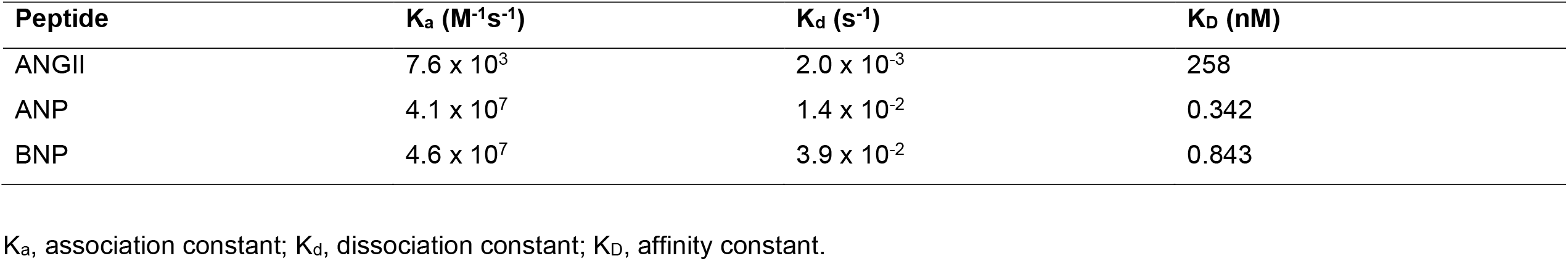
SPR Binding Kinetics for ANGII, ANP, and BNP to Human GC-A receptor.

| Peptide | $K_a$ ($M^{-1}s^{-1}$) | $K_d$ ($s^{-1}$) | $K_D$ (nM) |
| --- | --- | --- | --- |
| ANGII | $7.6 \times 10^3$ | $2.0 \times 10^{-3}$ | 258 |
| ANP | $4.1 \times 10^7$ | $1.4 \times 10^{-2}$ | 0.342 |
| BNP | $4.6 \times 10^7$ | $3.9 \times 10^{-2}$ | 0.843 |

$K_a$ , association constant; $K_d$ , dissociation constant; $K_D$ , affinity constant.

### Involvement of Protein Kinase C in the Interaction between ANGII/AT_1_ Signaling and GC-A

Finally, we explored the downstream mechanisms which mediate the interaction between the ANGII/AT_1_ signaling and the GC-A receptor. The protein kinase C (PKC) is a well-characterized downstream target of the ANGII/AT_1_ signaling, whose activation is mediated by phospholipase C (42,43). Importantly, the activation of PKC has been independently shown to desensitize the GC-A receptor through dephosphorylation (44,45). Through profiling of a series of different PKC isoforms (α, β, γ, δ, ε, and ζ), we found that protein expression of both PKCα and PKCε in the membrane fraction was substantially higher in HEK293/GC-A^+^/AT_1_^+^ cells compared to HEK293/GC-A^+^ cells (**Figure 5A**). Treatment of ANGII have no effect on PKC expression at protein level in both cell lines (**Figure 5A**). Therefore and to this end, we determined if PKC mediates the suppression effect of ANGII on GC-A. We characterized several known PKC small molecule modulators and found that treatment of Go6983 at 5 μM effectively inhibited PKC activity, represented by the phosphorylated status of pan-PKC substrates, in an mitogen-activated protein kinase (MAPK) independent manner (**Figure 5B**). Furthermore and in HEK293/GC-A^+^/AT_1_^+^ cells, treatment of both valsartan (1 μM), an AT_1_ receptor blocker, and Go6983 (5 μM) significantly rescued the suppression effect of ANGII on ANP-induced cGMP (**Figure 5C**). Altogether, these in vitro data additionally affirm the requirement of AT_1_ receptor and suggest the critical involvement of PKC in the underlying mechanisms of this RAAS-NPS crosstalk at a cellular level (**Figure 5D**).

**Figure 5.**
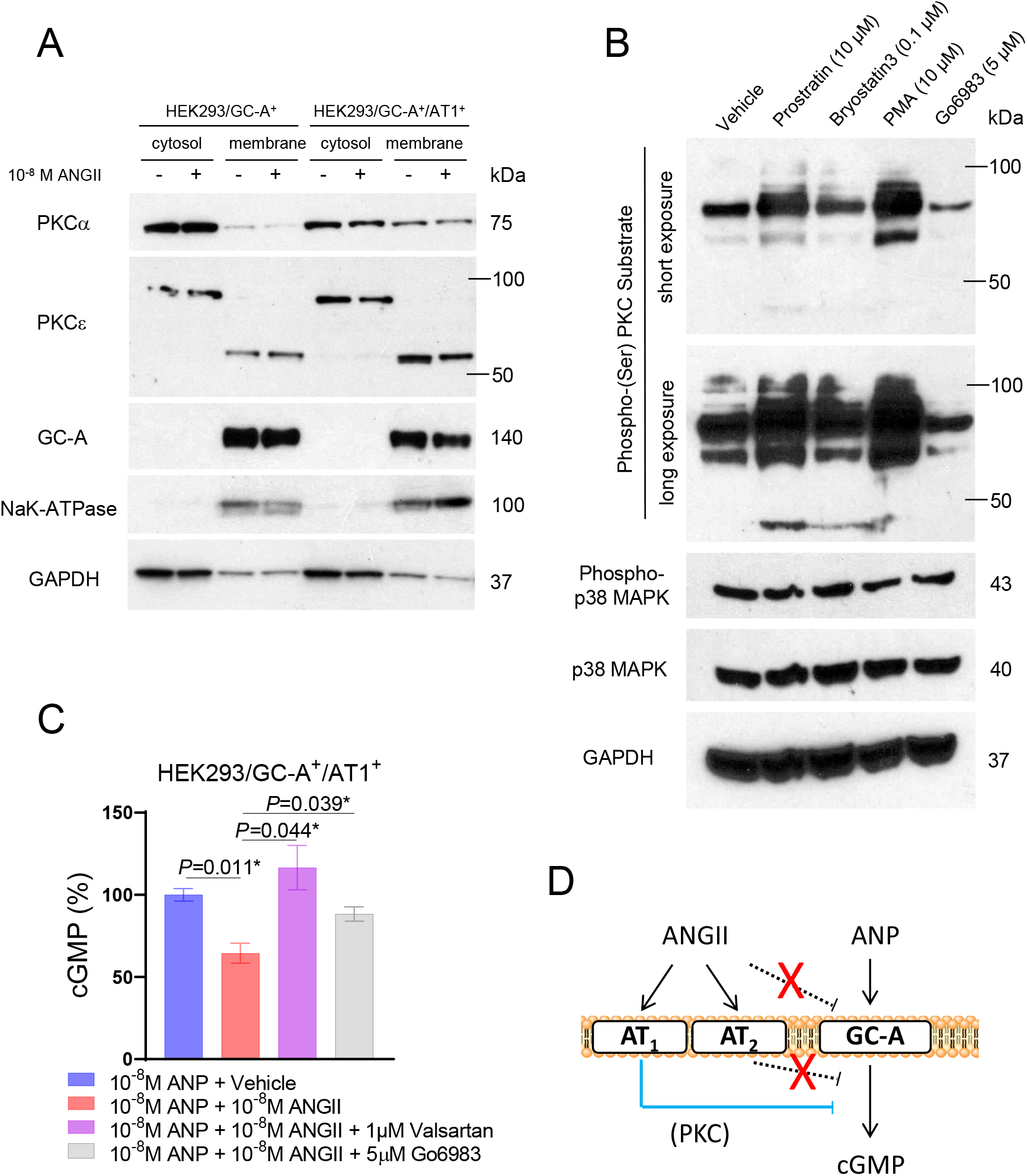
Involvement of PKC Signaling in the Crosstalk between ANGII/AT_1_ and GC-A. (A) Protein expression of different human PKC isoforms in the membrane and cytosol fractions of different HEK293 transfected cell lines (with or without treatment of ANGII). GC-A and NaK-ATPase serve as quality control for the separation of membrane and cytosol fractions. GAPDH serves as loading control. (B) Protein expression of phosphorylated PKC substrates, phosphorylated p38 MAPK, and p38 MAPK in HEK293/GC-A^+^/AT_1_^+^ cells treated with different PKC modulators. GAPDH serves as loading control. PMA, phorbol 12-myristate 13-acetate. (C) In vitro cGMP generation in HEK293/GC-A^+^/AT_1_^+^ in response to ANP (10^−8^ M) with or without ANGII (10^−8^ M) and valsartan (1 μM) or Go6983 (5 μM). Values of cGMP were normalized to vehicle group (blue) under ANP treatment. * indicates *P*<0.05, one-way ANOVA with Dunnett multiple comparisons test. (D) Hypothetical mechanism underlying the crosstalk between ANGII and GC-A derived from our in vitro studies.

## DISCUSSION

The present study was designed to advance the physiologic and mechanistic understanding of the pivotal crosstalk between the RAAS and the NPS. Specifically, we report findings from multiple approaches which all support that ANGII functions as a naturally occurring suppressor of the GC-A receptor. First, levels of circulating ANGII were found to influence the correlations between cGMP and endogenous GC-A ligands, ANP and BNP, in a cohort of healthy humans. Second, a co-infusion of ANGII with ANP in normal rats significantly attenuated cGMP production and the BP lowering response induced by ANP in vivo. Third, we demonstrated that the inhibitory effect of ANGII on GC-A receptor is through the AT_1_ receptor, but not through direct binding interaction with GC-A nor through the AT_2_ receptor in vitro. Fourth, we showed that PKC is an important mediator for the interaction between ANGII/AT_1_ signaling and the GC-A receptor. Altogether our findings support a fundamental but previously underappreciated concept that ANGII naturally suppresses GC-A and attenuates the generation of its second messenger cGMP via AT_1_ receptor and PKC.

While the counter-regulatory physiologic actions in regulating BP and cardiorenal homeostasis between the RAAS and NPS have long been recognized (24,25), the exact mechanisms underlying the crosstalk between these two key neurohormonal systems especially under physiological conditions remain largely unknown. Herein and most importantly, we extended our understanding by systematically investigating, for the first time to our knowledge, the neurohumoral profile of both systems in healthy humans. Our human study was conducted in a stringently defined healthy cohort who had no previous diagnosis of any CV or metabolic diseases and without any history of CV medication use. Of note, the circulating levels of renin and aldosterone were within the normal and physiologic range in all studied subjects. This healthy cohort offers a unique opportunity to decipher and interpret this crosstalk without confounding factors that is often associated with pathophysiologic stress and CV medications. Compared to reported levels in patients with CV disease (46-49), the circulating levels of ANGII, NPs, and cGMP were low in our healthy subjects, which indicates physiologic homeostatic activity for both RAAS and NPS. Nonetheless, a negative trend between ANGII with ANP or BNP was still observed, which suggest a counter-regulatory relationship between ANGII and NPs even under physiologic conditions. Comparatively, levels of CNP appears to more independent of ANGII in healthy subjects.

The pleiotropic beneficial actions associated with the NPS on both heart and kidneys are primarily conferred through the generation of its second messenger cGMP (5,50), thus making this pathway an attractive therapeutic target as well as a readout of target engagement by the NPs. Indeed, cGMP modulation has emerged as one of the most promising mechanistic-based therapeutic strategies for CV drug discovery in recent years (51,52). In humans, circulating levels of cGMP are considered as a signal of NPS activation, and circulating levels of cGMP were reproducibly found to be elevated in association with increased levels of NPs in patients with HF (53-55). Here, through regression analysis, we also found circulating cGMP can be statistically predicted by the circulating levels of any single NP alone in a base model including age, sex, BMI, and eGFR in our healthy cohort. Interestingly, our data further demonstrates that adding ANGII levels and the interaction term between ANGII and NP into the model improved the predictive accuracy of cGMP particularly as it related to ANP and BNP, but less with CNP.

Prompted by the regression analysis, the current study further investigated neurohormonal profiles in subjects subgrouped by the circulating levels of ANGII. The most important observation is that both ANP and BNP have positive correlations with cGMP but only in those subjects with relatively low ANGII (below the median level). In subjects with relatively high ANGII (higher than the median level), the expected correlation between cGMP and either ANP or BNP disappeared. While this finding should be validated in larger cohorts, our study provides the first human evidence capturing the potential suppression of ANGII on NP associated cGMP. The fact that a higher level of ANGII even within the physiologic range disrupts the ANP/BNP/cGMP signaling further demonstrats this cross-system interaction is fundamental and naturally exists. Once again, a similar pattern was not observed with CNP following the same analytical approach, which together with the data mentioned earlier lead us to speculate that the interaction between ANGII and the NPS is predominantly on the GC-A axis rather than the GC-B axis. Nonetheless, studies may still be warranted to investigate if CNP and the GC-B receptor, as well as GC-A and its ligands, has any interaction with the RAAS components during human CV disease.

We and others are actively pursuing innovative GC-A enhancing therapies whose biological actions go beyond endogenous NPs as therapeutic drugs for CV and other diseases (56-59). Sangaralingham and colleagues (37) recently reported the discovery of a small molecule, GC-A positive allosteric modulator, which enhances the affinity of NPs to the GC-A receptor and augments cGMP generation. Along these lines, it remains unclear if any endogenous or exogenous negative modulator of GC-A also exists. Given the counter-regulatory actions between the RAAS and NPS, it is tempting to speculate that some RAAS components may serve as negative modulators of the NP receptors. Indeed, based on previous reports exploring ANGII effects on the NPS at multiple levels (32,33,60-63), ANGII may serve as a negative modulator of GC-A peptides from a drug development perspective. Herein we confirmed this hypothesis building on our in vitro studies in normal rats in vivo and found that co-infusion of a low dose of ANGII with ANP significantly attenuated ANP-induced cGMP levels in the plasma and urine, which was also accompanied by an attenuation of ANP-induced BP reduction. The dose of ANGII applied in this study was also optimized to rule out the complication of ANGII mediated cGMP alteration, as ANGII has been previously reported to stimulate the activity of PDEs for cGMP degradation (61,64). Our data suggest that an in vivo environment of high ANGII is suboptimal for the therapeutic actions of GC-A targeted NPs, further raising the possibility to enhance the favorable pharmacological effects of NPs with a combinatory use of RAAS inhibitors. Indeed, a blunted effect of ANP or neprilysin inhibitor was also reported in patients (65,66), and in a large animal (67) and rodent model (60) with chronic HF, a pathophysiologic condition with high ANGII activity. Future investigations are warranted to determine if the natural suppression effect of ANGII on ANP-induced cGMP we observed herein is the primary mechanism underlying those observed blunted effects of NPs in CV disease.

The importance of the crosstalk between the RAAS and NPS is strengthened, in part, by the recognition of the known complementary localization of functional receptors for both systems in a variety of organs including brain, adrenal gland, vasculature, heart, and kidneys (24). At the receptor level, two possible mechanisms of the observed desensitization of GC-A by ANGII were explored in current study. First, ANGII may directly bind to GC-A receptor and serve as a negative allosteric modulator for GC-A activity on cGMP production. Second, ANGII may counteract the activity of GC-A via triggering downstream signaling of its own receptor(s). Our in vitro data supports the latter hypothesis, as we observed no attenuation of ANP-induced cGMP in the presence of ANGII in HEK293 cells overexpressing GC-A alone, and the binding affinity between ANGII and GC-A was found to be low, unlike ANP and BNP which had strong binding to GC-A that is consistent with a previous study (37). While this finding appears contradictory to the early studies which demonstrated the inhibition of ANGII on ANP-induced cGMP in glomerular mesangial cells (33) and aortic smooth muscle cells (32), we further found this interaction was only recapitulated in HEK293 cells when AT_1_ was co-overexpressed together with GC-A. This mechanism was specific to AT_1_, as overexpressing AT_2_ together with GC-A resulted in no change in ANP-induced cGMP generation in our study. Notably, the requirement for the presence of AT_1_ receptor was additionally validated by the rescuing of ANP-induced cGMP in the presence of valsartan, a commonly used AT_1_ blocker in clinical practice.

Compared to previously used rat primary cells with expression of multiple receptors (32-34), the baseline expression of the GC-A, AT_1_, and AT_2_ were found to be minimal in parental HEK293 cells. Herein through side-by-side comparison among three engineered cells, we provide clear evidence that the crosstalk between ANGII and GC-A on cGMP generation is dependent on the AT_1_ receptor. Our data thus far further support an important involvement of PKC, a downstream kinase of the AT_1_ receptor, in the crosstalk between ANGII/AT_1_ and GC-A. While the exact mechanism deserve further invetigations, this finding appears to be in line with the previous knowledge that the phosphorylation status of GC-A is critical for its activity (44,68). Future studies are warranted to determine if PKC-associated phosphorylation on GC-A is the primarily mechanism which can explain the observed ANGII suppression on ANP induced cGMP herein. Meanwhile, the influence of ANGII on GC-A can be multi-dimensional and beyond the regulation of GC-A functional activity. For instance, ANGII has also been recently reported to affect GC-A expression at mRNA and protein levels (62,63,69). Whether and how these mechanisms may also be involved in the crosstalk between ANGII/AT_1_ and GC-A deserve additional studies and emphasis especially in different disease states.

## STUDY LIMITATIONS

Our study is a combination of statistical approach in human data, in vivo physiologic validation, and in vitro mechanistic exploration. One limitation is that it is challenging to build a definite linkage between experimental and human observations. Thus, different approaches were designed in the current study in parallel to provide novel insights from multiple perspectives to facilitate translation of preclinical studies to the human environment. The entire study was demonstrated in an acute setting and mostly under physiologic conditions, and future studies need to extend findings here to chronic pathophysiologic states such as hypertension and heart failure.

## CONCLUSIONS

In summary, our studies provide new knowledge which support ANGII as a functional suppressor of GC-A receptor activity in humans, in vivo, and in vitro. These findings further advance our understanding of the critical crosstalk between the ANGII/AT_1_ and NP/GC-A/cGMP signaling pathways. We also conclude that our findings highlight the promising therapeutic avenue to design strategies targeting both pathways simultaneously to optimize the beneficial effects seen with GC-A/cGMP activation.

## NONSTANDARD ABBREVIATIONS AND ACRONYMS

ACE: angiotensin converting enzyme
ARBs: AT_1_ receptor blockers
ANGII: angiotensin II
AT_1_: ANGII type 1 receptor
BP: blood pressure
cGMP: 3’, 5’ cyclic guanosine monophosphate
CV: cardiovascular
GC-A: particulate guanylyl cyclase A
NPS: natriuretic peptide system
NP: natriuretic peptide
PKC: protein kinase C
RAAS: renin angiotensin aldosterone system
S/V: sacubitril/valsartan

## ACKNOWLEDGEMENTS

The authors thank Christopher G. Scott for his statistical support and advice.

## DATA AVAILABILITY

All data related to the findings in current study are available upon request in reasonable time and form.

## FIGURE LEGENDS

**Supplemental Table 1.**
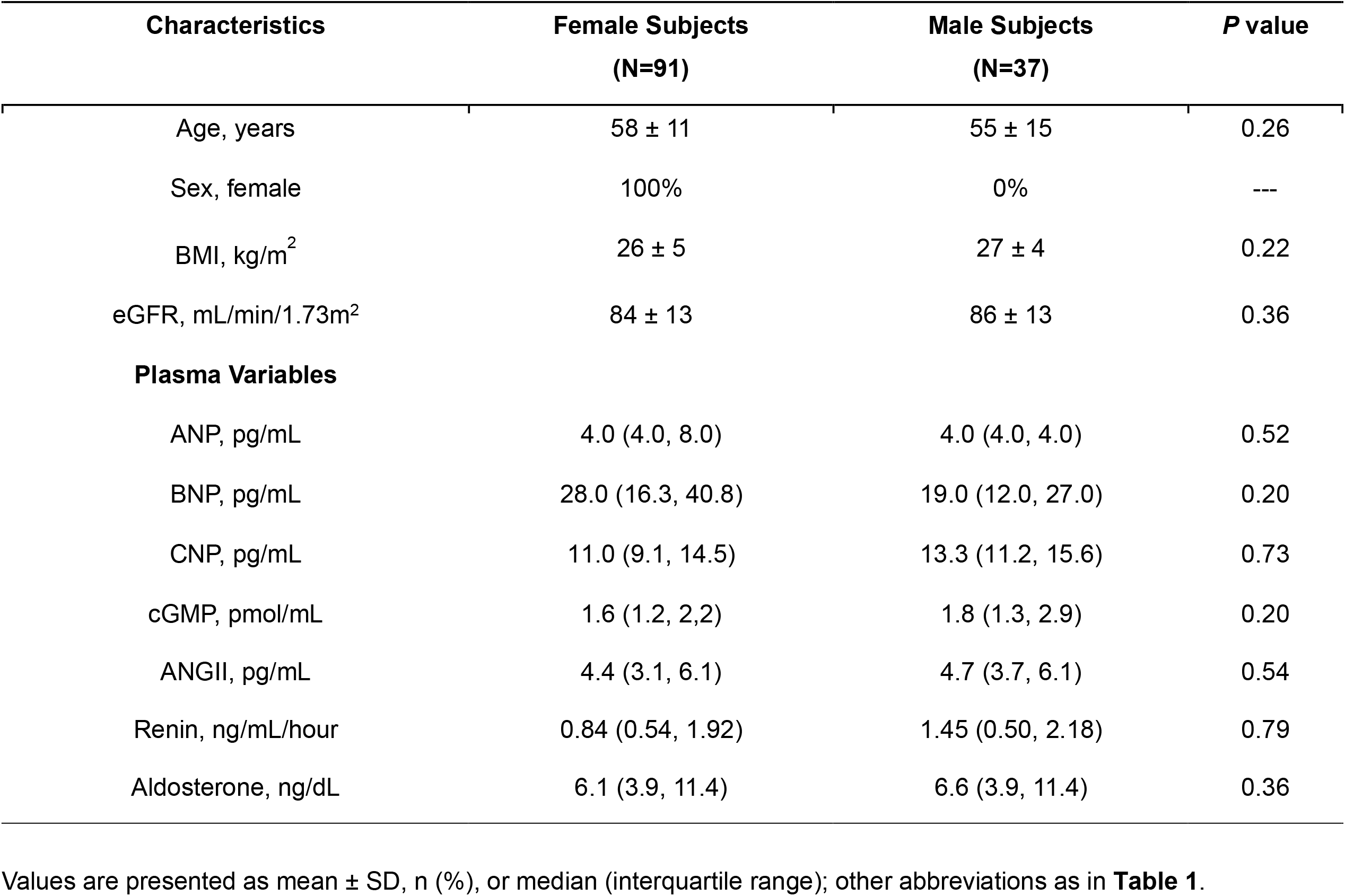
Baseline Characteristics of Healthy Subjects Subgrouped by Sex.

**Supplemental Table 2.** Univariable Analysis of Plasma ANGII in Healthy Subjects.

| N = 128 |  | Log<sub>10</sub>(Plasma ANGII) |  |
| --- | --- | --- | --- |
| Predictor Variable | β (SE) | r | P Value |
| Age | -0.003 (0.002) | -0.13 | 0.15 |
| Sex, female | -0.02 (0.03) | -0.03 | 0.58 |
| BMI | 0.0004 (0.0005) | 0.002 | 0.94 |
| eGFR | 0.001 (0.002) | 0.07 | 0.43 |
| Log<sub>10</sub>(Plasma ANP) <sup>#</sup> | -0.13 (0.09) | -0.24 | 0.12 |
| Log<sub>10</sub>(Plasma BNP) | -0.14 (0.08) | -0.16 | 0.07 |
| Log<sub>10</sub>(Plasma CNP) | -0.03 (0.13) | -0.02 | 0.81 |
| Log<sub>10</sub>(Plasma cGMP) | -0.18 (0.10) | -0.14 | 0.11 |

Log<sub>10</sub> transformation was performed on plasma ANP, BNP, CNP, cGMP, and ANGII for approximate normality of distribution.

**Supplemental Figure 1.**
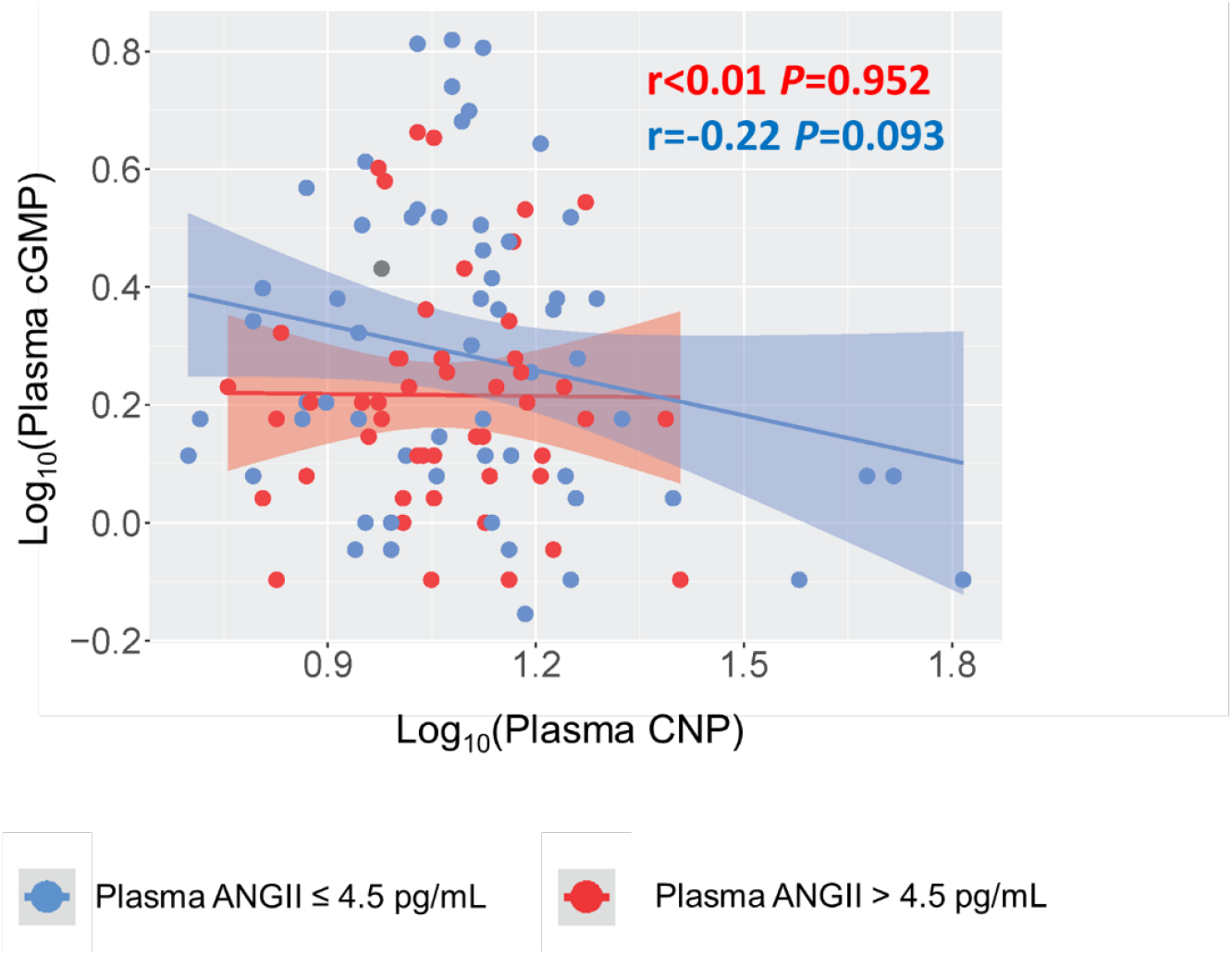
Correlations between cGMP and CNP in Healthy Subjects Grouped by ANGII Levels. Linear correlation between plasma CNP and plasma cGMP. Each dot represents one healthy subject. Blue indicates the subjects with plasma ANGII ≤ 4.5 pg/mL; Red indicates the subjects with plasma ANGII > 4.5 pg/mL. Pearson correlation coefficient r and *P* value were shown for each constructed linear regression model.

**Supplemental Figure 2.**
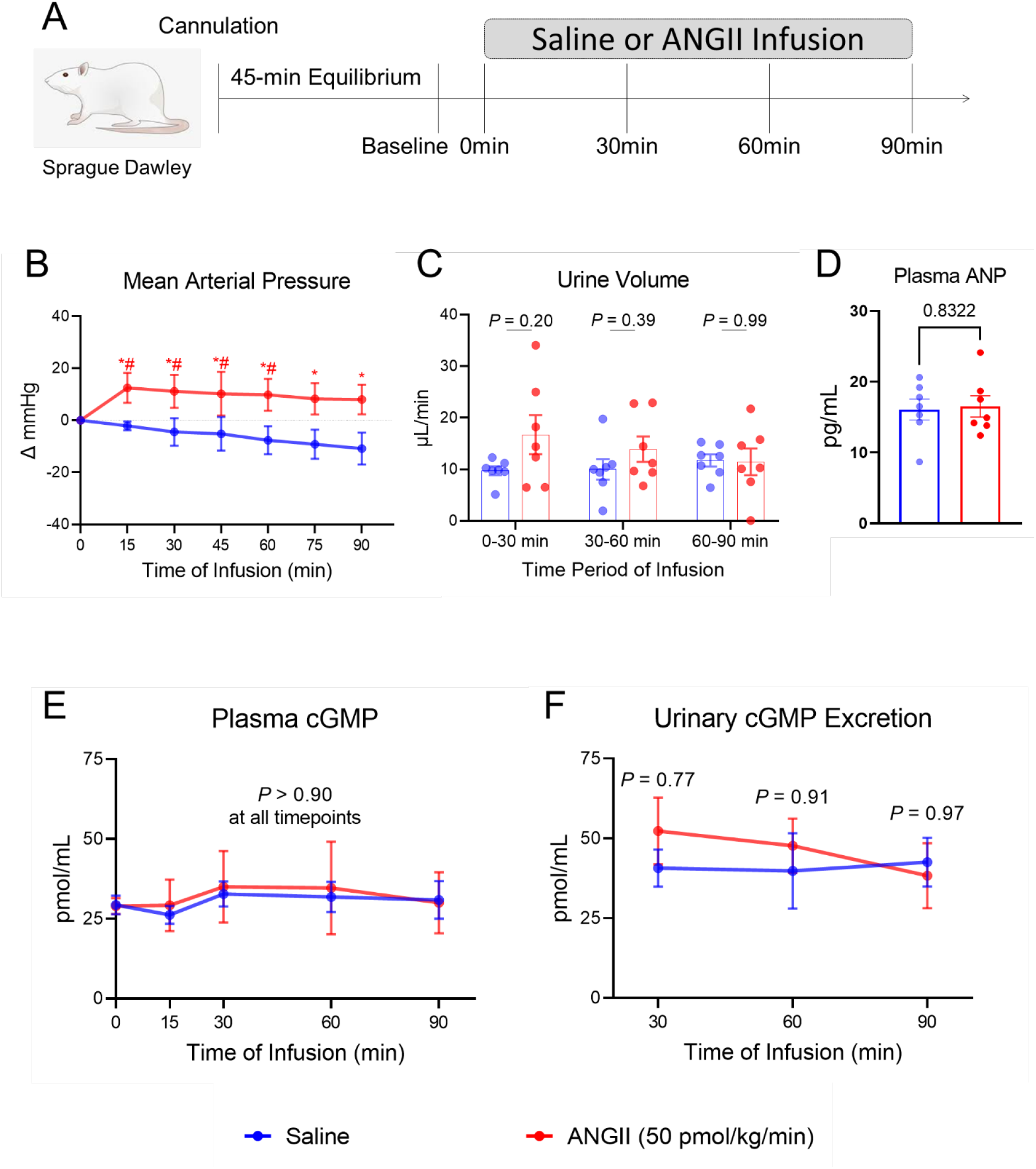
ANGII Effect on Endogenous cGMP In Vivo. (A) The acute protocol study design to evaluate the effect of ANGII infusion in normal rats. (B) Mean arterial pressure. * indicates *P*<0.05 compared to saline group; # indicates *P*<0.05 compared to corresponding baseline. (C) Urinary output represented by urine volume collected at every 30-min interval during continuous infusion. (D) Plasma ANP at 90 mins after saline or ANGII infusion. (E) Plasma cGMP, no statistical difference was found between two groups or compared to baseline at all timepoints. (F) Urinary cGMP excretion. no statistical difference was found between two groups or compared to baseline at all timepoints. N=7 in saline group, N=7 in ANGII group.

**Supplemental Figure 3.**
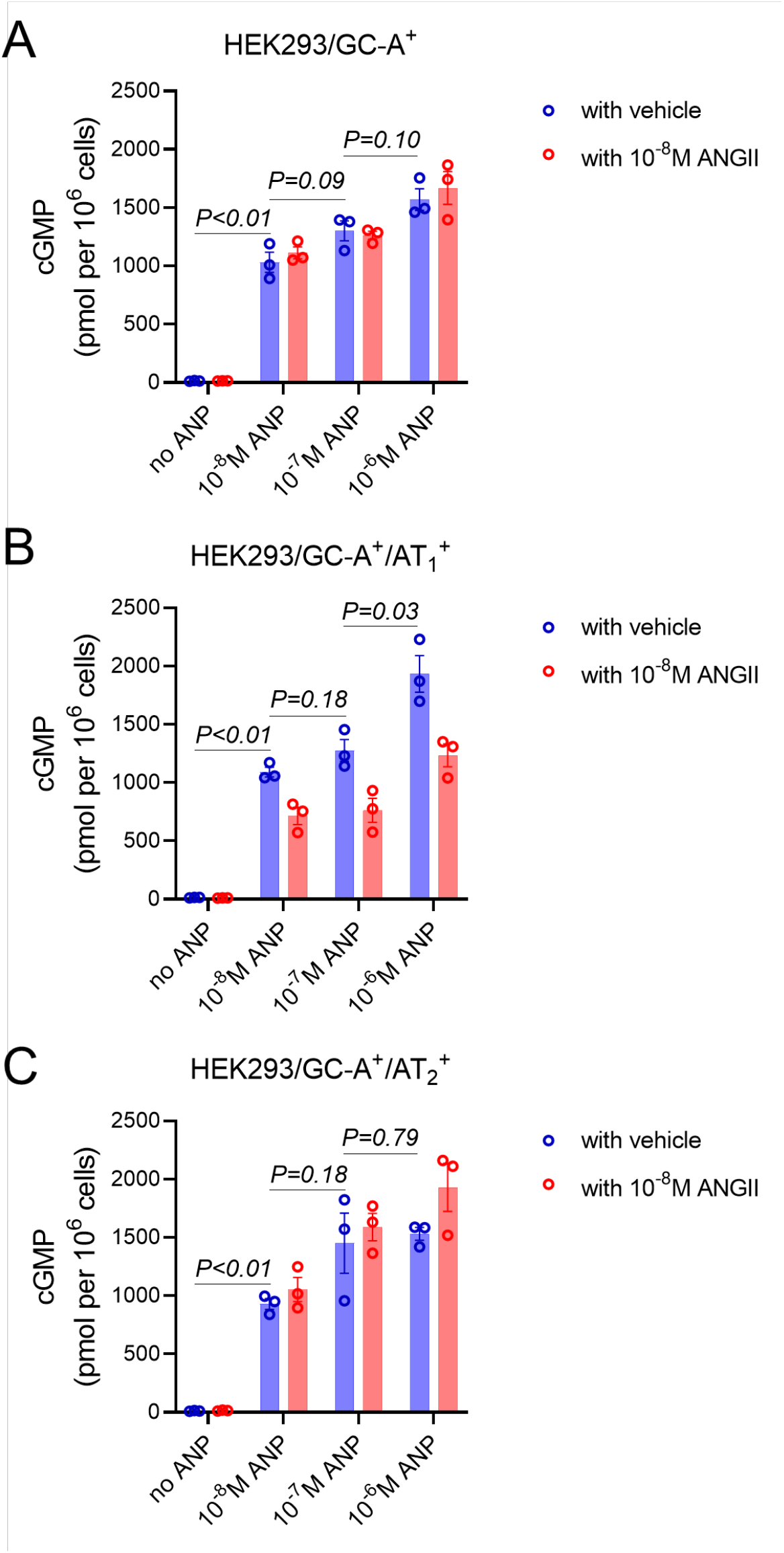
In Vitro cGMP Generation in HEK293 Transfected Cells. Absolute values of cGMP generation in different HEK293 transfected cells in response to ANP with or without ANGII (10^−8^ M). The effect of vehicle alone or ANGII alone in corresponding HEK293 transfected cells were labeled as “no ANP” as negative control. Values of cGMP were quantified in pmol per 10^6^ cells. Each dot represents one biological replicate. Shown *P* values are comparisons between different doses of ANP without ANGII (blue bars) using two-side unpaired t test.

**Supplemental Figure 4.**
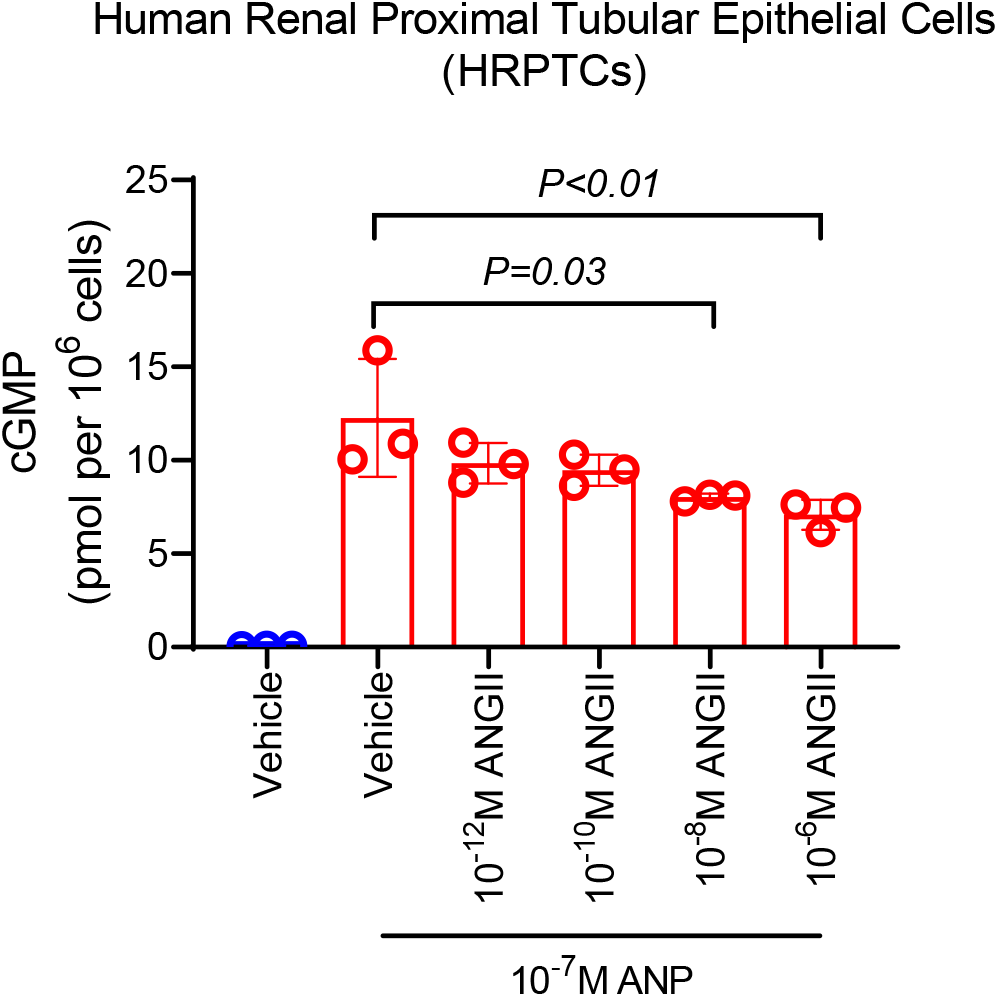
In Vitro cGMP Generation in Human Renal Proximal Tubular Epithelial Cells. Absolute values of cGMP generation in human renal proximal tubular epithelial cells (HRPTCs) in response to ANP (10^−7^M) with or without different concentrations of ANGII (10^−12^ to 10^−6^ M). Values of cGMP were quantified in pmol per 10^6^ cells. Each dot represents one biological replicate. Only significant *P* values (<0.05) are shown in the figure using one-way ANOVA following by multiple comparison.

